# Experimentally reduced insulin/IGF-1 signalling in adulthood extends lifespan of parents and improves Darwinian fitness of their offspring

**DOI:** 10.1101/405019

**Authors:** Martin I. Lind, Sanjana Ravindran, Zuzana Sekajova, Hanne Carlsson, Andrea Hinas, Alexei A. Maklakov

## Abstract

Classical theory maintains that ageing evolves via energy trade-offs between reproduction and survival leading to accumulation of unrepaired cellular damage with age. In contrast, the emerging new theory postulates that ageing evolves because of deleterious late-life hyper-function of reproduction-promoting genes leading to excessive biosynthesis in late-life. The hyper-function theory uniquely predicts that optimizing nutrient-sensing molecular signalling in adulthood can simultaneously postpone ageing and increase Darwinian fitness. Here we show that reducing evolutionarily conserved insulin/IGF-1 nutrient-sensing signalling via *daf-2* RNA interference (RNAi) fulfils this prediction in *Caenorhabditis elegans* nematodes. Long-lived *daf-2* RNAi parents showed normal fecundity as self-fertilizing hermaphrodites and improved late-life reproduction when mated to males. Remarkably, the offspring of *daf-2* RNAi parents had higher Darwinian fitness across three different genotypes. Thus, reduced nutrient-sensing signalling in adulthood improves both parental longevity and offspring fitness supporting the emerging view that sub-optimal gene expression in late-life lies at the heart of ageing.

**Impact Statement:** Understanding mechanisms underpinning ageing is fundamental to improving quality of life in an increasingly long-lived society. Recent breakthroughs have challenged the long-standing paradigm that the energy trade-off between reproduction and somatic maintenance causes organismal senescence via slow accumulation of unrepaired cellular damage with age. The emerging new theory of ageing provides a conceptually novel framework by proposing that ageing is a direct consequence of physiological processes optimized for early-life function, such as growth and early-life reproduction, that are running ‘too high’ (i.e. at hyperfunction) in late adulthood. Contrary to the classic view based on damage accumulation, the hyperfunction theory proposes that suboptimal gene expression in late-life causes ageing via excessive biosynthesis. Thus, the hyperfunction theory uniquely predicts that longevity and Darwinian fitness can be simultaneously increased by reducing unnecessarily high levels of nutrient-sensing signalling in adulthood. Here we show that reducing evolutionarily conserved nutrient-sensing signalling pathway fulfils this prediction in *Caenorhabditis elegans* nematodes. We found that downregulation of the insulin/IGF-1 signalling in adult *C. elegans* nematodes not only improves longevity but, most intriguingly, increases fitness of the resulting offspring in the next generation. We found support for increase in offspring fitness across different genetic backgrounds. Our findings contradict the theoretical conjecture that energy trade-offs between growth, reproduction and longevity is the universal cause of senescence and provide strong experimental support for the emerging hyperfunction theory of ageing.

## Introduction

Understanding mechanisms underpinning healthy ageing is fundamental to improving quality of life in an increasingly long-lived society. The long-standing paradigm postulates that energy trade-offs between reproduction and somatic maintenance underlie organismal ageing (Kirkwood 1977, Kirkwood and Austad 2000, Kirkwood 2017). This theory is supported by a large number of studies in different taxa that reported a negative correlation between reproduction and survival (reviewed in Hughes and Reynolds 2005, Partridge et al. 2005, Edward and Chapman 2011, Flatt 2011, Nussey et al. 2013). However, the discoveries of environmental interventions that dramatically increase healthy lifespan in model organisms without the cost of reduced reproduction have challenged the current paradigm and suggested that our understanding of the evolution of ageing is incomplete (Dillin et al. 2002, Kenyon 2005, Antebi 2013, Gems and Partridge 2013, Maklakov and Immler 2016, Flatt and Partridge 2018). Specifically, experimental downregulation of nutrient-sensing insulin/IGF-like (IIS) signalling pathway that governs biosynthesis in response to nutrient availability can achieve increased longevity without a concomitant decrease in reproduction in model organisms (Dillin et al. 2002, Kenyon 2010, Kenyon 2011).

Since cost-free lifespan extension contradicts the traditional view of how ageing evolves, several studies investigated the fitness consequences of reduced IIS signalling (Gems et al. 1998, Walker et al. 2000, Jenkins et al. 2004, Savory et al. 2014, Maklakov et al. 2017). Indeed, mutations that reduce nutrient-sensing signalling during the whole life, as well as environmental interventions aimed at mimicking the mutational effect, often have detrimental pleiotropic effects on key life-history traits, such as development, growth rate, body size and early-life reproduction resulting in reduced Darwinian fitness even if total reproduction is unaffected (Gems et al. 1998, Briga and Verhulst 2015). The first longevity mutant discovered in *C. elegans, age-1*, is a good example because increased longevity, stress resistance and late-life reproduction come at a cost of reduced early-life reproduction and total individual fitness (Maklakov et al. 2017). Moreover, a recent literature survey suggests that all classic longevity-extending mutations across taxa from worms to flies to mice detrimentally affect life-history traits resulting in reduced fitness (Briga and Verhulst 2015). Similarly, experimental evolution studies showed that when longevity and fecundity are increased simultaneously through selection, the organisms pay the price in slow development and delayed sexual maturation, again resulting in reduced individual fitness (Lind et al. 2017). These results support the theoretical conjecture that genes with antagonistically pleiotropic effects between early-life and late-life fitness play an important role in the evolution of ageing (Williams 1957). However, the mechanisms of antagonistic pleiotropy (AP) remain elusive. The leading hypothesis, the “disposable soma” theory of ageing (DS) suggests that ageing results from competitive energy allocation between somatic maintenance and reproduction (Kirkwood 1977, Kirkwood and Holliday 1979, Kirkwood and Austad 2000). Indeed, increased reproductive performance in early-life correlates with reduced survival and/or reduced performance in late-life in natural populations (Gustafsson and Part 1990, Boonekamp et al. 2014, Lemaitre et al. 2015) and in laboratory experiments (Rose 1984, Charlesworth 1993, Partridge et al. 1999, but see Chen and Maklakov 2012, Kimber and Chippindale 2013, Chen and Maklakov 2014, Curtsinger 2019).

However, this hypothesis suffered several setbacks in recent years, with many empirical studies challenging the importance of energy trade-offs in organismal senescence (reviewed in Flatt 2011, Kenyon 2011, Antebi 2013, Gems and Partridge 2013, Maklakov and Immler 2016, Flatt and Partridge 2018). Instead, several authors proposed that ageing can result from molecular signalling networks being optimized for development, growth and early-life reproduction rather than for late-life reproduction and longevity (Blagosklonny 2010, Kenyon 2010, Antebi 2013, Gems and Partridge 2013, Ezcurra et al. 2018). For example, the hyperfunction theory maintains that ageing is driven by excessive nutrient-sensing molecular signalling in adulthood, which results in cellular hypertrophy leading to age-related pathologies (Blagosklonny 2006, 2010, Ezcurra et al. 2018). These ideas can be traced back to the original AP theory by George Williams, who suggested that the same physiological processes that are beneficial for fitness early in life can become detrimental for organismal fitness with age because of the reduced strength of natural selection on late-life function (Williams 1957).

Williams’s AP theory (Williams 1957) provides the population genetic framework for the evolution of ageing via two different physiological routes: energy trade-offs (the “disposable soma” theory of ageing) or functional trade-offs (e.g. “hyperfunction” theory of ageing) (Fig. 1). While both of these physiological theories rely on the same underlying principle, they make uniquely distinct predictions with respect to reproduction costs of longevity. The “disposable soma” theory maintains that organismal senescence is caused by slow accumulation of unrepaired cellular damage with age because insufficient energy resources are allocated to repair as organisms are maximising fitness rather than longevity (Kirkwood 1977, Kirkwood 2017). Therefore, the “disposable soma” theory predicts that increased allocation of resources to somatic maintenance will increase longevity at the cost of reduced resources available to current growth and reproduction. On the contrary, the functional trade-off theory suggests that longevity is compromised by suboptimal physiology in adulthood because, as discussed first by Williams (1957), selection is not strong enough to fully optimize the age-specific expression of an allele whose effects are strongly beneficial in early-life (e.g. during development) and slightly detrimental in late-life (e.g. during adulthood). Consequently, the functional trade-off theory predicts that experimental optimization of physiology in adulthood can increase longevity without any cost to reproduction, or even simultaneously increase longevity and reproduction.

**Fig. 1.**
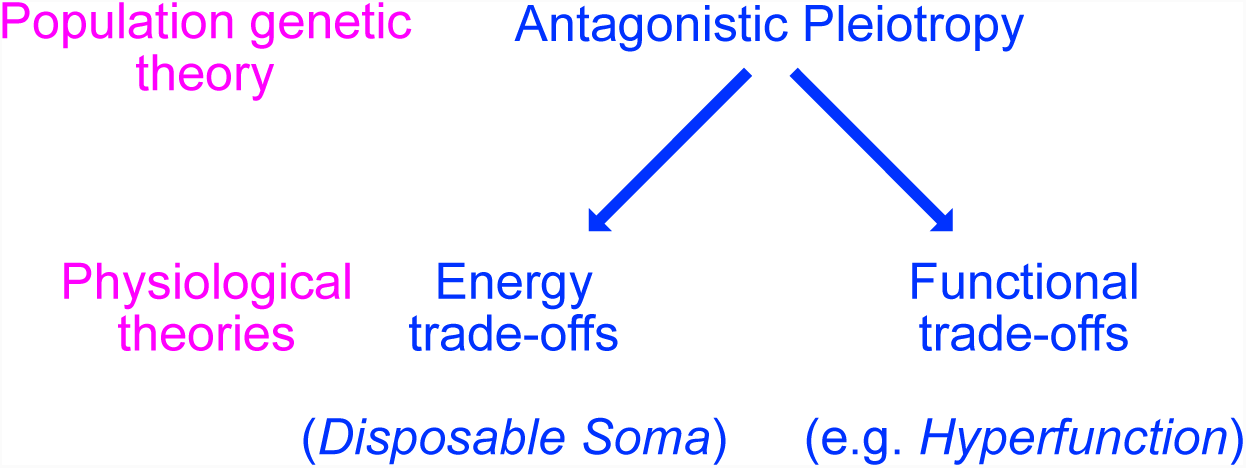
Theoretical framework. The relationship between population genetic theory of ageing (“antagonistic pleiotropy”, Williams 1957) and physiological theories of ageing based on either energy trade-offs (“disposable soma” (Kirkwood 1977) or functional trade-offs (e.g. “hyperfunction” (Blagosklonny 2006)). Note that Williams (1957) was the first to provide an abstract example of a functional trade-off between early-life and late-life allelic effects on organismal physiology. Recently, Blagosklonny (2006, 2010) put forward a “hyperfunction” hypothesis that specifically links suboptimal levels of nutrient-sensing signalling to excessive biosynthesis (hence, hyperfunction) leading to cellular and organismal senescence.

Because the main cost of longevity appears to be associated with reduced early-life function, it seems plausible that age-specific modification of gene expression can potentially circumvent this problem. In their landmark study, Dillin et al. (2002) used age-specific RNA interference (RNAi) approach to knock down *daf-2* gene expression in *C. elegans* nematodes across the life cycle of the worms. While early-life feeding with bacteria expressing *daf-2* double-stranded (ds) RNA resulted in reduced early-life reproduction, there was no detrimental effect of *daf-2* RNAi in adult worms, which enjoyed two-fold lifespan extension without any cost to reproduction (Dillin et al. 2002). This study provides the strongest support to date for the hypothesis that ageing results from molecular nutrient-sensing signalling that is optimized for early-life function but is suboptimal for late-life function.

Nevertheless, while this study provided a powerful example for the cost-free lifespan extension, it is possible that certain fitness costs have been overlooked. One possibility is that fecundity costs become apparent only in mated hermaphrodites. In nature, *C. elegans* live in populations with small (∼0.3%) yet appreciable number of males living among self-fertilising hermaphrodites with sometimes high levels of outcrossing (Sivasundar and Hey 2005), and mating, as well as mere presence of male-derived pheromones, has pronounced effects of the life-history of hermaphrodites (Maures et al. 2013, Shi and Murphy 2014, Aprison and Ruvinsky 2016). While it is certainly interesting to consider how male-derived effects affect resource allocation in hermaphrodites, it is unlikely that putative trade-off that is only visible to selection in such rare circumstances can shape the evolution of ageing in this species. Perhaps more importantly, it is possible that the fitness of the offspring and, therefore, Darwinian fitness of the parents are compromised. The trade-off between offspring number and quality is well known from a number of study systems (Stearns 1992), and is a potential explanation for the apparent lack of fitness costs in the previous studies (Maklakov and Immler 2016). Alternatively, longevity and Darwinian fitness can be simultaneously increased by reducing unnecessarily high levels of nutrient-sensing signalling in adulthood. To distinguish between these possibilities, we need to understand how reduction in nutrient-sensing signalling in adulthood affects longevity, offspring number and offspring quality. Here we show that *daf-2* RNAi in adult *C. elegans* results in increased offspring fitness across three genetic backgrounds. We discuss these findings in the light of the emerging new theories of ageing and suggest that they support the hypothesis that functional trade-offs between early-life fitness and late-life fitness shape the evolution of ageing.

## Materials and Methods

### Strains

We used the *Caenorhabditis elegans* strains Bristol N2 wild-type (Brenner, Genetics 1974), as well as the mutants *ppw-1(pk2505)* and *rrf-1(pk1417)*, obtained from Caenorhabditis Genetics Center (CGC, Missouri, USA).

### Maintenance

Before each assay, worms were recovered from freezing and synchronised by bleaching for two generations to remove any freezing effects. The nematode populations were maintained at 20°C and 60% relative humidity in an environmental test chamber. For regular maintenance, the worms were kept on NGM agar supplemented with the antibiotics streptomycin, kanamycin and nystatin (following Lionaki and Tavernarakis 2013), seeded with the antibiotic-resistant *E. coli* strain OP50-1 (pUC4K).

### Outline of the study

The study was run in three separate experiments. In the first experiment, we investigated lifespan and reproduction of mated and unmated N2 hermaphrodites reared from sexual maturity onwards on *daf-2* RNAi or empty vector (EV, control) plates. For logistic reasons, this experiment was conducted in two blocks for mated worms and one block for unmated worms. In the second experiment, we investigated the lifespan and egg size of unmated N2, *rrf-1(pk1417)* and *ppw-1(pk2505)* hermaphrodites on raised from sexual maturity onwards on *daf-2* RNAi or EV plates. In the third experiment, we collected one egg from each parent at their second day of adulthood (from *daf-2* RNAi and EV treatments) and investigated the lifespan and reproduction of these offspring on control plates. Because different experiments differed in setup time, daily reproduction values (and calculations based upon these, such as λ_ind_) are only meaningful for comparison between treatments within each experiment.

### RNAi

RNase-III deficient, IPTG-inducible HT115 *Escherichia coli* bacteria with empty plasmid vector (L4440) was used as control (Timmons et al. 2001) and the same HT115 bacteria with *daf-2* RNAi construct from the Vidal library was used as RNAi treatment. RNAi treatment started from sexual maturity, and continued until the death of the individual. During the experiments, worms were maintained on 35 mm NGM agar plates (supplemented with 1 mM IPTG and 50 μg/ml ampicillin) seeded with 0.1 ml L4440 empty vector control or *daf-2* bacteria grown in LB supplemented with 50 μg/ml ampicillin for 16-20 hours and seeded (incubated) on the NGM agar plates again for 24 hours (following. Hinas et al. 2012).

### Lifespan Assays

Lifespan assays were set up for all treatment combinations described above. In the lifespan assays, the individual age-synchronised L4 worms were placed on separate 35 mm plates and the plates were checked daily to record any instances of death. The surviving worms were moved to new plates daily until their death. Fertile worms, which showed odd developmental characteristics and low offspring numbers (<36 offspring), were excluded from the final analysis (3 mated control worms and 7 mated *daf-2* worms).

### Reproduction assays

Offspring production was scored in the reproduction assays using the same worms as those scored for lifespan, except for the parental N2, *ppw-1* and *rrf-1* worms in the second experiment, where only lifespan was recorded. Unmated individual hermaphrodites were moved to new plates daily and scored for offspring produced 2.5 days later. In the “mated” treatment, two male *C. elegans* (from the initial sample population of N2 strain) were placed on a plate with a single hermaphrodite for two hours every day to allow time for mating. Offspring production was scored 2.5 days later, as in the “unmated” treatment.

### Egg size assays

Egg size was measured in N2, *ppw-1* and *rrf-1* strains (unmated hermpahrodites) growing on either *daf-2* RNAi or empty vector (EV) plates. Two days after maturation, worms were placed individually on new plates and observed continually during five hours for the presence of newly laid eggs, of which the first two eggs were collected. Eggs were picked immediately after laying and placed under a Leica M165C microscope set on 12x magnification; photos were taken using a Lumenera Infinity 2-6C digital microscope camera. Egg size was analysed from photos using *ImageJ* (https://imagej.nih.gov/ij/). Only eggs laid during gastrulation stage (the normal developmental stage at egg laying) were included in the analyses.

### Statistical analyses

Survival was analysed for each experiment in Cox proportional hazard models in *R 3.3.3*. Mated (EV: n=72, *daf-2* n=68) and unmated (n=25 per treatment) individuals were analysed separately, as they were run in different blocks. Unmated individuals were analysed using the *coxph* function in the package *survival*, with *daf-2* RNAi treatment as a fixed factor. For mated individuals, we used the *coxme* package in order to fit block as a random effect, in addition to the fixed effect of RNAi treatment. In the second experiment (n=25 per treatment), in addition to RNAi treatment, we also fitted the fixed factor strain (N2, *ppw-1, rrf-1*) and its interaction with treatment using the *coxph* function in the *survival* package.

Reproduction was analysed as total reproduction as well as rate-sensitive individual fitness λ_ind_, which encompasses the timing and number of offspring (Brommer et al. 2002, Lind et al. 2016). λ_ind_ is estimated by solving the Euler-Lotka equation for each individual using the *lambda* function in the *popbio* package and is analogous to the intrinsic rate of population growth (Stearns 1992). For all unmated worms (n=25 per treatment), we estimated the fixed effect of treatment (*daf-2* RNAi or empty vector). For offspring of the three mutants (n=25 per treatment), we also estimated the fixed effect or strain, using linear models. For the mated worms (EV: n=72, *daf-2* n=68), we also estimated the random effect of block, in addition to RNAi treatment. These models were implemented as mixed effect models using the *lme4* package in *R 3.3.3*, and chi-square tests of fixed effects were performed using the *car* package. Egg size was analysed in a mixed effect model in *lme4*, treating strain and RNAi treatment as crossed fixed effects, and parent ID as well as block as random effects. We obtained the following n: N2 on EV: 56, N2 on *daf-2*: 54, *ppw-1* on EV: 44, *ppw-1* on *daf-2*: 42, *rrf-1* on EV: 59, *rrf-1* on *daf-2*: 42.

## Results

First off, we confirmed that *daf-2* RNAi significantly extended the lifespan of unmated N2 wild-type hermaphrodite worms (censoring matricide: z = −4.94, df = 1, p <0.001, Fig. 2A; including matricide as dead: z = −4.97, df = 1, p <0.001), as expected from previous studies (Dillin et al. 2002). In addition, for mated N2, *daf-2* RNAi extended lifespan when matricide was censored (z = −2.42, df = 1, p = 0.016, Fig. 2B) but not if matricidal worms were included as dead (z = 0.16, df = 1, p = 0.87) because of an increase in matricide in the late reproducing mated *daf-2* RNAi N2.

**Fig. 2.**
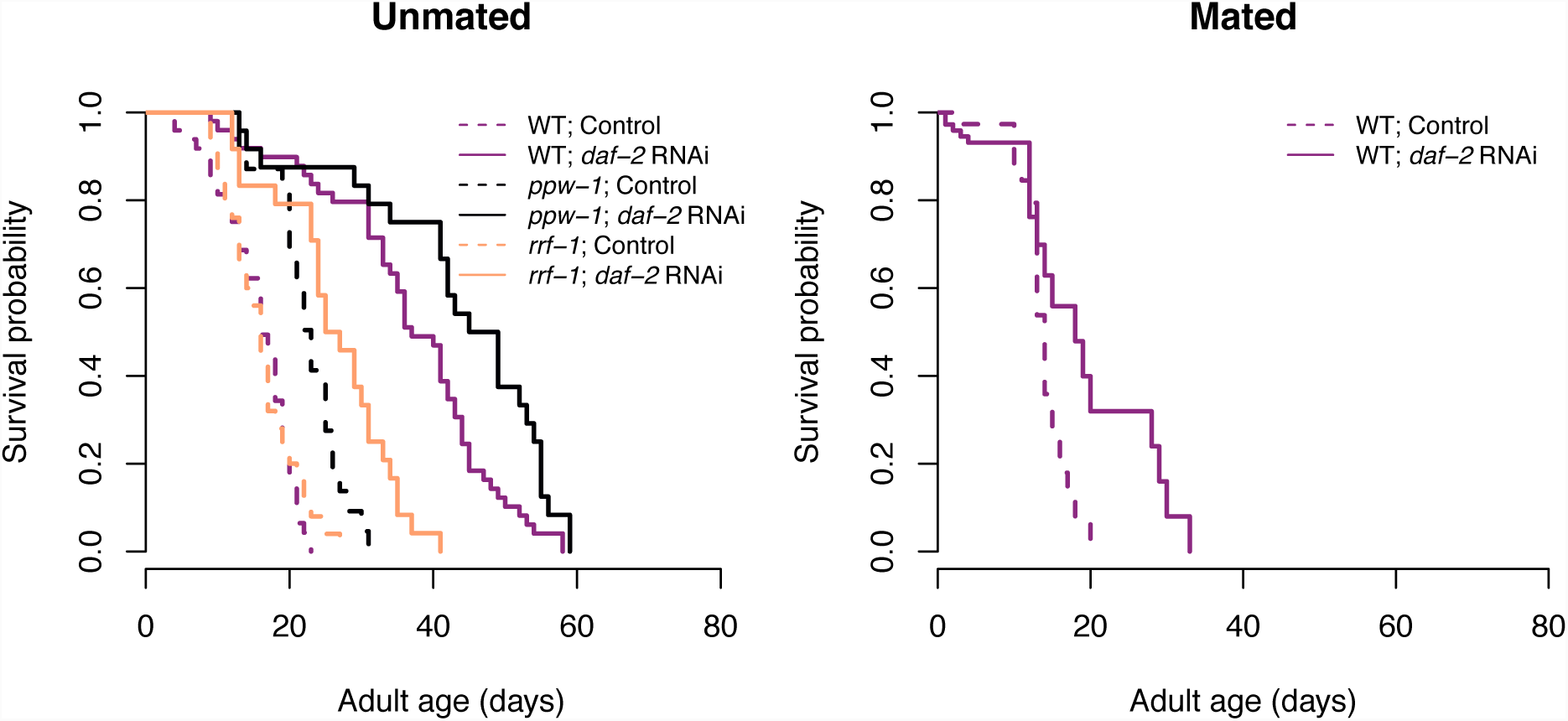
The effect of *daf-2* RNAi on lifespan. Survival probability for (**A**) unmated or (**B**) mated N2 wild-type (purple), *ppw-1* (black) and *rrf-1* (orange) mutants, treated with either *daf-2* RNAi (solid lines, filled symbols) or control empty vector (broken lines, open symbols) from adulthood onwards.

We did not find any effect of *daf-2* RNAi on total reproduction (unmated: F = 0.32, df = 1, p = 0.58; mated: χ^2^ = 1.11, df = 1, p = 0.29) or individual fitness λ_ind_ (unmated: F = 0.30, df = 1, p = 0.59; mated: χ^2^ = 0.43, df = 1, p = 0.51) for neither unmated nor mated N2 (Table 1, Fig. 3). However, *daf-2* RNAi had a positive effect on late (day 5+) reproduction for mated hermaphrodites (χ^2^ = 24.76, df = 1, p <0.001, Fig. 3B).

**Table 1.** The effect of daf-2 RNAi on reproduction. Total reproduction and individual fitness (λ_ind_) for unmated and mated *C. elegans* N2 wild-type treated with either empty vector (Control) or *daf-2* RNAi from adulthood onwards. All values expressed as mean ± SE.

| RNAi treatment | Total reproduction | | Fitness ($\lambda_{\text{ind}}$) | |
| --- | --- | --- | --- | --- |
|  | unmated | mated | unmated | mated |
| Control | 311.0 ± 7.0 | 595.7 ± 24.3 | 4.66 ± 0.03 | 4.47 ± 0.05 |
| *daf-2* RNAi | 317.4 ± 8.9 | 630.5 ± 22.2 | 4.63 ± 0.05 | 4.50 ± 0.05 |

**Fig. 3.**
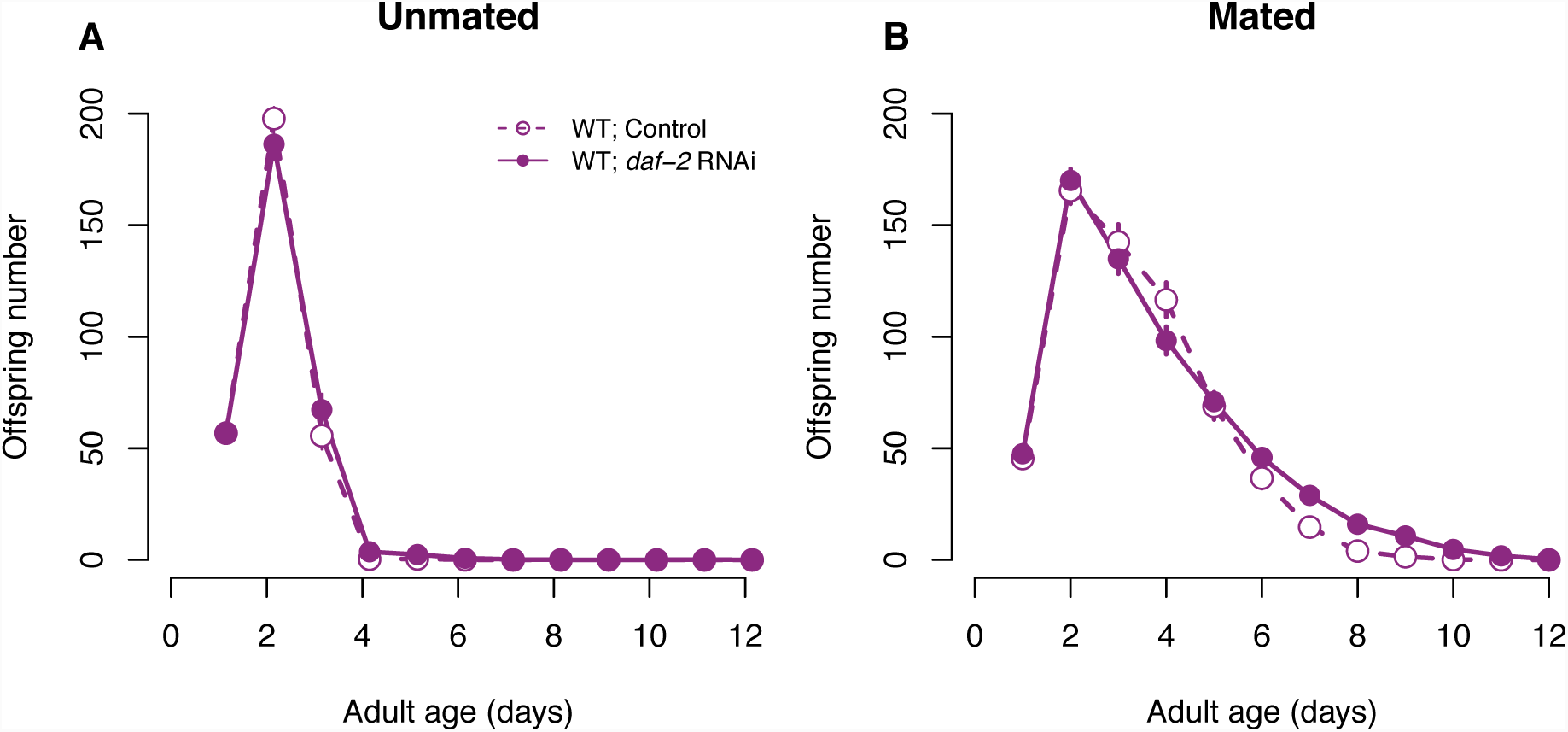
The effect of *daf-2* RNAi on reproduction. Daily offspring number for (**A**) unmated or (**B**) mated N2 wild-type worms, treated with either *daf-2* RNAi (solid lines, filled symbols) or control empty vector (broken lines, open symbols) from adulthood onwards. Symbols represent mean ± SE.

In a second experiment, using unmated hermaphrodites only, we investigated the effect of *daf-2* RNAi on parent lifespan and offspring lifespan and reproduction across three genetic backgrounds (N2 wild-type and the mutants *ppw-1* and *rrf-1*, that are deficient for germline and somatic RNAi, respectively). Parental treatment with *daf-2* RNAi increased lifespan across all genetic backgrounds, both when matricide was censored (treatment: χ^2^ = 90.39, df = 1, p <0.001; strain: χ^2^ = 21.8, df = 2, p <0.001; treatment × strain: χ^2^ = 10.46, df = 2, p = 0.005, Fig. 2A) and included as dead (treatment: χ^2^ = 85.25, df = 1, p <0.001; strain: χ^2^ = 20.45, df = 2, p <0.001; treatment × strain: χ^2^ = 9.43, df = 2, p = 0.009). In accordance with previously published research (Hibshman et al. 2016), parental *daf-2* RNAi increased egg size (treatment: χ^2^ = 5.11, df = 1, p =0.024; strain: χ^2^ = 13.89, df = 2, p <0.001; treatment × strain: χ^2^ = 2.68, df = 2, p = 0.262, Fig. 4). However, we found that the effect was most pronounced in N2 wildtype worms, and relatively weak in both somatic and germline *daf-2* knockdown (see Fig. 4), suggesting that *daf-2* knockdown in both somatic and reproductive tissues is required to maximize the effect on egg size.

**Fig. 4.**
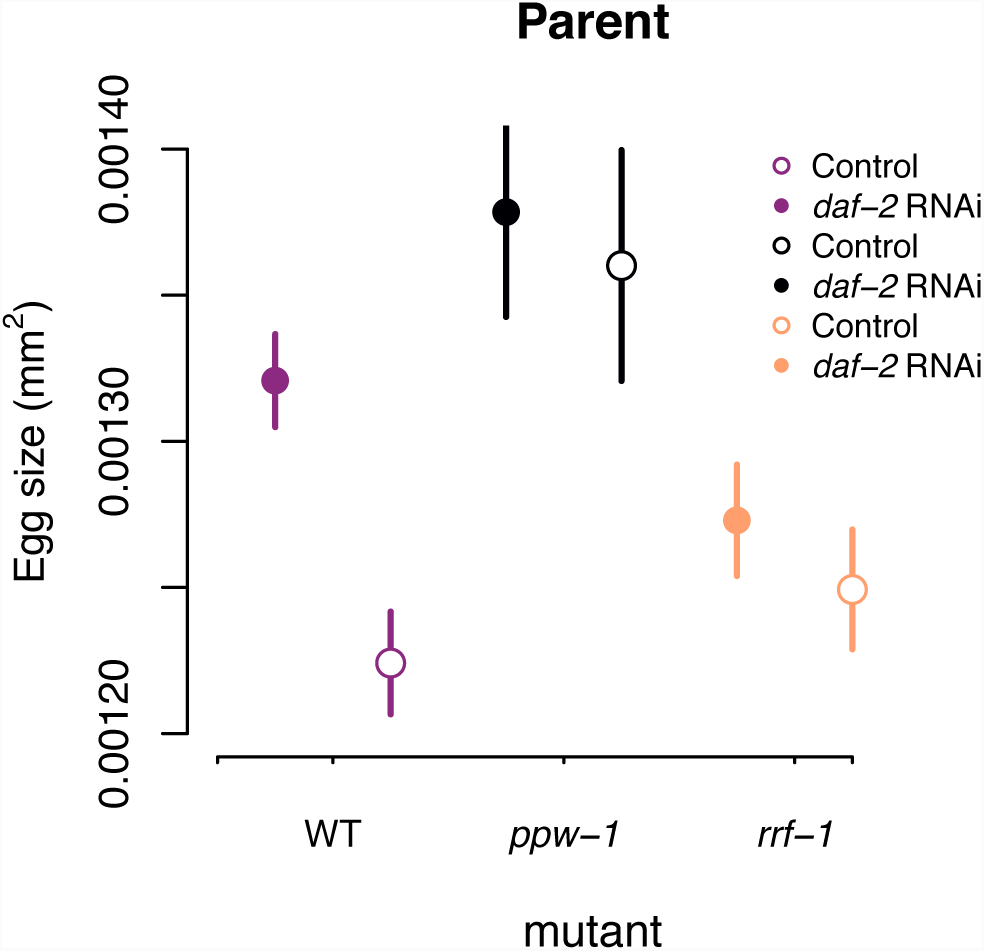
The effect of *daf-2* RNAi on egg size. Egg size of unmated, parental worms. N2 wild-type (purple), *ppw-1* (black) and *rrf-1* (orange) mutants, treated with either *daf-2* RNAi (solid lines, filled symbols) or control empty vector (broken lines, open symbols) from adulthood onwards. Symbols represent mean ± SE.

Parental *daf-2* RNAi treatment did not, however, influence the lifespan of their offspring, neither when matricidal worms were censored (treatment: χ^2^ = 0.04, df = 1, p = 0.85; strain: χ^2^ = 24.2, df = 2, p <0.001; treatment × strain: χ^2^ = 0.61, df = 2, p = 0.74, Fig. 5A) nor when included as dead (treatment: χ^2^ = 0.01, df = 1, p = 0.92; strain: χ^2^ = 21.8, df = 2, p <0.001; treatment × strain: χ^2^ = 0.48, df = 2, p = 0.79).

**Fig. 5.**
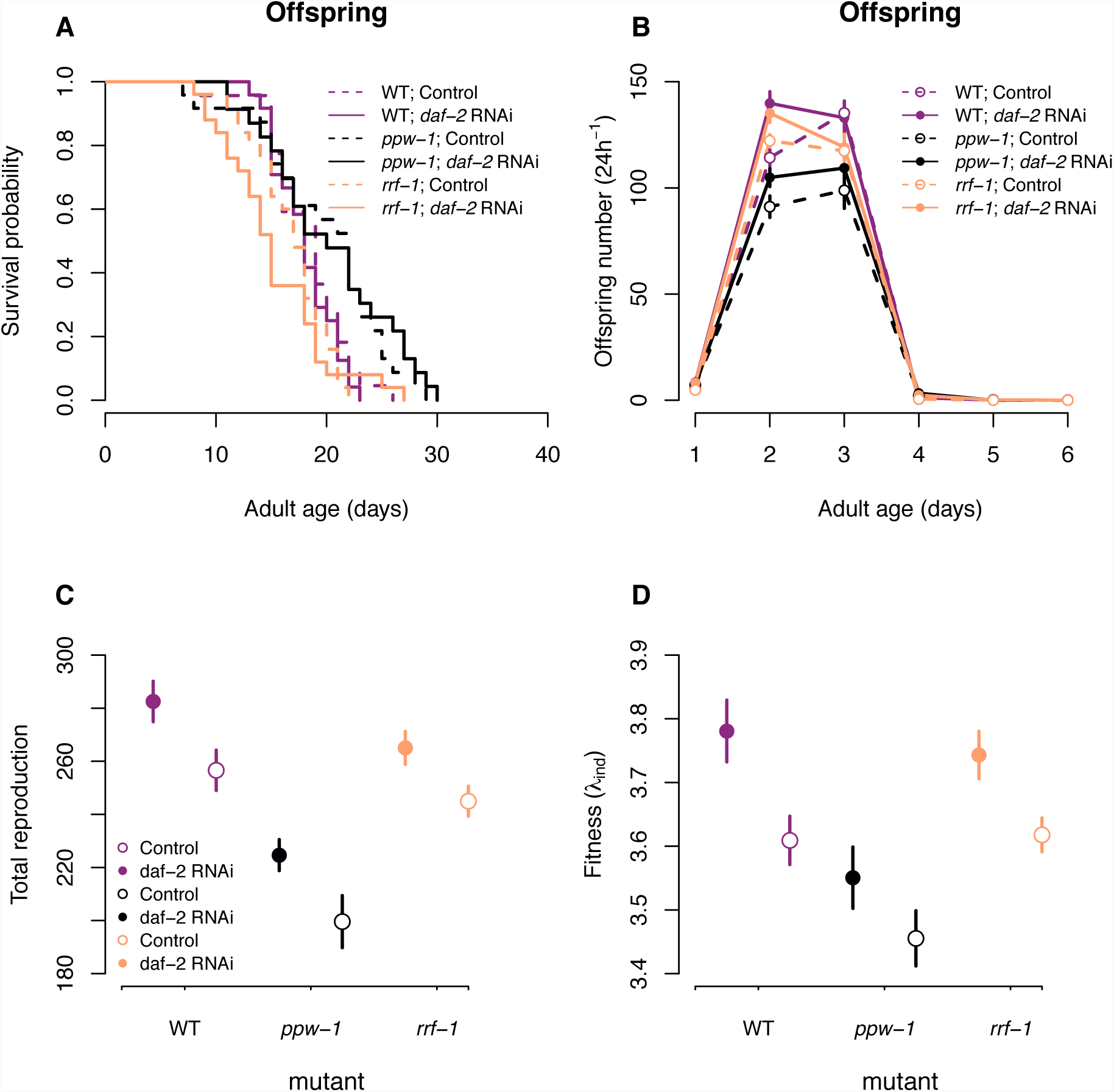
The effect of parental *daf-2* RNAi on offspring survival and reproduction. Offspring worms, unmated, on control (empty vector) plates from parents exposed to *daf-2* RNAi or control treatment. (**A**) Survival probability, (**B**) daily offspring number, (**C**) Total reproduction and (**D**) individual fitness (λ_ind_) of offspring (on control plates) from parents either exposed to *daf-2* RNAi (solid lines, filled symbols) or control empty vector (broken lines, open symbols). The colors reflect N2 wild-type (purple), *ppw-1* (black) and *rrf-1* (orange) mutants. Symbols represent mean ± SE.

In contrast, parental *daf-2* RNAi treatment significantly increased offspring total reproduction (treatment: F = 15.9, df = 1, p <0.001; strain: F = 33.7, df = 2, p <0.001; treatment × strain: F = 0.09, df = 2, p = 0.91, Fig. 5B-C) and individual fitness λ_ind_ (treatment: F = 11.8, df = 1, p <0.001; strain: F = 13.1, df = 2, p <0.001; treatment × strain: F = 0.18, df = 2, p = 0.84, Fig. 5D) across all genetic backgrounds. Importantly, there was no correspondence between the effect of parental *daf-2* RNAi on egg size (see above) and offspring total reproduction / individual fitness, suggesting that factors beyond the amount of resources in the egg contribute to increased fitness of offspring of *daf-2* RNAi parents.

## Discussion

The “disposable soma” theory of ageing postulates that senescence results from competitive energy allocation between key life-history traits, such as growth, reproduction and somatic maintenance (Kirkwood 1977, Kirkwood 2017). This theory predicts that genetic and environmental manipulations that increase energy allocation to growth and somatic maintenance will result in detrimental effects to reproduction. This is why the findings by Dillin et al. (2002), which suggested that adult-only downregulation of insulin/IGF-1 by *daf-2* RNAi can substantially increase lifespan without any detrimental effect to reproduction, were subsequently scrutinized in an attempt to find the hidden costs of longevity (Jenkins et al. 2004, Partridge et al. 2005). Nonetheless, both the original findings (Dillin et al. 2002) and our results here, suggest that adult-only *daf-2* RNAi can more than double longevity without any negative effect on reproduction. Moreover, when supplied with sperm from males, *daf-2* RNAi-treated parents have improved fecundity in late-life. However, the key question that we asked in this study was whether treatments that improve parental performance have positive or negative effects on their offspring. The trade-off between offspring number and offspring quality is a well known concept in life-history evolution (Stearns 1992) but is rarely considered in biogerontological research (reviewed in Maklakov and Immler 2016). Germline maintenance is costly (Sniegowski et al. 2000, Agrawal and Wang 2008, Maklakov and Immler 2016, Berger et al. 2017), and increased investment into somatic maintenance can, in theory, result in increased mutation rate and reduced fitness of progeny.

Alternatively, it is possible that instead of energy trade-offs, the evolution of senescence is governed by functional trade-offs. Functional trade-offs can occur because the physiological requirements of a young organism can differ substantially from those of a mature one (Williams 1957). In his classic 1957 paper, George Williams (Williams 1957) described a hypothetical example of a mutation that positively affects bone calcification in a developing young organism but increases calcification of the connective tissues of arteries in a mature one with detrimental consequences. More recently, it has been suggested that nutrient sensing IIS/TOR molecular signalling pathways that govern growth and development result in excessive biosynthesis in late-life leading to different pathologies and increased mortality (Blagosklonny 2006, 2010, Gems and Partridge 2013, Ezcurra et al. 2018). These proximate explanations rest on the fundamental assumption that the strength of natural selection declines with age because of environmental mortality from a range of biotic and abiotic hazards (e.g. predation, pathogens, competition, starvation) (Williams 1957). Because of such environmental mortality, optimizing development, growth and instantaneous reproduction may be more important for organismal fitness that optimizing future survival and reproduction (Williams 1957, Hamilton 1966). Thus, progressively weakening selection in adulthood may result in suboptimal levels of IIS/TOR signalling leading to pathology and senescence (Ezcurra et al. 2018). However, unlike the classic energy trade-off theory, the functional trade-off hypothesis predicts that it should be possible to modify adult physiology to improve both longevity and fitness.

Here we found that reduced insulin/IGF-1 signalling in adult worms not only improved longevity, but also increased reproduction and Darwinian fitness of the resulting offspring in three different genetic backgrounds. This result contradicts the hypothesis that improved longevity and postponed ageing of *daf-2* RNAi parents comes at the cost of offspring fitness. Instead, our findings are in line with the hypothesis that suboptimal levels of nutrient-sensing signalling in adult life accelerate ageing, curtail lifespan and reduce individual fitness. This result was not caused by direct inheritance of *daf-2* RNAi, since we did not recover the lifespan extension effect of *daf-2* knockdown in these offspring. Because *daf-2* is essential for successful development and growth of a young worm (Dillin et al. 2002), these results suggest that wildtype *C. elegans* nematodes trade-off improved pre-adult performance for reduced offspring quality. Such trade-offs are at heart of AP theory, but are usually interpreted as evidence for energy-based trade-offs. Our results clearly demonstrate that this is not the case, and that adulthood-only *daf-2* RNAi increases offspring fitness. In summary, our findings suggest that selection on expression of *daf-2* in adulthood is not sufficiently strong in nature. We predict that such effects may be very common, and suggest that future studies should aim to quantify the fitness consequences of experimental manipulation of age-specific gene expression across a broad range of taxa. Such approach will allow us to estimate the relative importance of energy trade-offs versus functional trade-offs in the evolution of ageing across the tree of life.

Because previous research found that both dietary restriction and reduction in insulin-like signalling by *daf-2* RNAi knockdown increased embryo size in *C. elegans* nematodes (Hibshman et al. 2016), we replicated these results to test whether increased fitness of adult progeny results from increased resource allocation to eggs by *daf-2* RNAi mothers. While *daf-2* knockdown increased egg size to a different degree in N2, *ppw-1* and *rrf-1* strains, there was no correlation between the effect of parental *daf-2* RNAi on egg size and offspring reproductive performance. We provisionally conclude that increased egg size under reduced maternal insulin-like signalling can contribute to increased offspring fitness, but it is likely not the sole source of variation in this trait. Recent work has identified oocyte-specific IIS targets that are different from soma-specific IIS targets suggesting that IIS signalling regulates reproduction and longevity through different mechanisms (Templeman et al. 2018). In the future, it will be interesting to test for individual fitness of offspring produced via genetic manipulation of oocyte-specific targets of IIS signalling pathway. In recent years, there has been a vigorous debate regarding whether mechanistic understanding of life-history trade-offs is necessary to advance life-history theory (Flatt and Heyland 2011, Stearns 2011a, b). Here we used the mechanistic approach to separate between two conceptually different evolutionary theories of ageing – energy trade-offs and functional trade-offs – in an empirical study. We argue that unification between the conceptual approach and the mechanistic understanding may often prove fruitful in this regard.

## Author contributions

MIL and AAM designed the study, with the aid of AH. SR, ZS, MIL and HC collected the data, MIL analysed the data, MIL and AAM drafted the manuscript. All authors contributed to the revision of the manuscript.

## Acknowledgements

This work was supported by ERC Consolidator Grant 724909 GermlineAgeingSoma to AAM and Swedish Research Council VR Grant 2016-05195 to MIL. The authors declare no conflicts of interest.

## Data accessibility

Upon acceptance, the data will be archived at Dryad.

